# LoL-align: sensitive and fast probabilistic protein structure alignment

**DOI:** 10.1101/2025.11.24.690091

**Authors:** Lasse Reifenrath, Michel van Kempen, Gyuri Kim, Soo Hyun Kim, Mohammadreza Radnezhad, Milot Mirdita, Martin Steinegger, Johannes Söding

## Abstract

The ubiquitous availability of protein structures permits replacing sequence alignment with more accurate and sensitive structure alignment algorithms. LoL-align maximizes a <u>lo</u>cal log-odds score for proteins to be homologous, given their intra-protein *C*_*α*_ – *C*_*α*_ distances. LoL-align is markedly more sensitive in detecting remote homologs than TMalign and DALI on single- and multi-domain benchmarks and achieves better alignment quality at 5-20× their speed. LoL-align is fully integrated into Foldseek.

---

Advances in protein structure prediction now yield models with near-experimental accuracy [1, 2, 3, 4]. Today’s databases contain hundreds of millions of predicted structures [5, 2], which transforms large-scale protein analyses from sequence-to structure-based approaches and creates the need for accurate and efficient structural alignment tools.

Accurate structure aligners should detect remote homology with high sensitivity and specificity. They should produce accurate residue-level alignments, also for full-length multidomain proteins. The ability to detect local similarities and tolerance to conformational variability are essential. Speed is equally critical, as millions of structures need to be compared in large-scale analyses.

Existing methods meet these requirements only partially. TM-align [6] and the fast, GPU-enabled GTalign superimpose proteins in 3D space, offering accurate alignments for rigid, single-domain structures but limited tolerance for conformational flexibility. Dali [7, 8], which compares intramolecular distance patterns, remains among the most sensitive methods [9], but its runtime precludes large-scale use. Foldseek [10] compresses structures into a 20-state 3Di alphabet and applies MMseqs2-style searches [11], achieving 10^4^–10^5^-fold speedups, but with reduced sensitivity and alignment quality. Several deep learning–based approaches have been developed [12], although they may struggle when applied to proteins outside their training distribution.

Here we present LoL-align, a distance-based algorithm that iteratively optimizes our novel Local-Log-odds scoring function. The LoL-score quantifies the evidence for a pair of residues to be homologous given their *C*_*α*_–*C*_*α*_ distances and sequence distances within a 20Å neighborhood. LoL-align achieves higher sensitivity than Dali at 20 times higher speed. It is tolerant to conformational flexibility and produces more accurate alignments on average than TM-align and Dali. Integrated into Foldseek, LoL-align can refine top hits after prefiltering, enabling large-scale structural comparisons with high accuracy.

With LoL-align, we also present the first algorithm for SIMD-vectorized Forward–Backward probabilistic sequence alignment [13], which achieved substantial performance gains through three optimizations: (i) SIMD-accelerated dynamic programming for parallel processing on modern CPU architectures; (ii) a rescaling scheme to prevent numerical over- and underflow, which allows us to perform calculations in linear space without costly logarithmic transformations; (iii) a small memory footprint by overloading dynamic programming matrices.

LoL-align first runs the Forward–Backward algorithm using as match scores the sum of 3Di and amino acid substitution scores (Fig. 1). The top ten aligned 7-mers with the highest sum of posterior alignment probabilities are extended along the diagonal of the dynamic programming matrix, and the three top-scoring 7-mer seed anchors are retained for further refinement. For each seed anchor, the alignment is iteratively refined in three steps per iteration. First, a LoL-score matrix is computed using the current anchor pair residues. Second, posterior alignment probabilities are computed with the Forward–Backward algorithm. Third, residues exceeding a probability threshold are added to the set of anchor residues. In this way, the anchor residue set expands locally in 3D space until convergence is achieved (Fig. 1).

**Figure 1.**
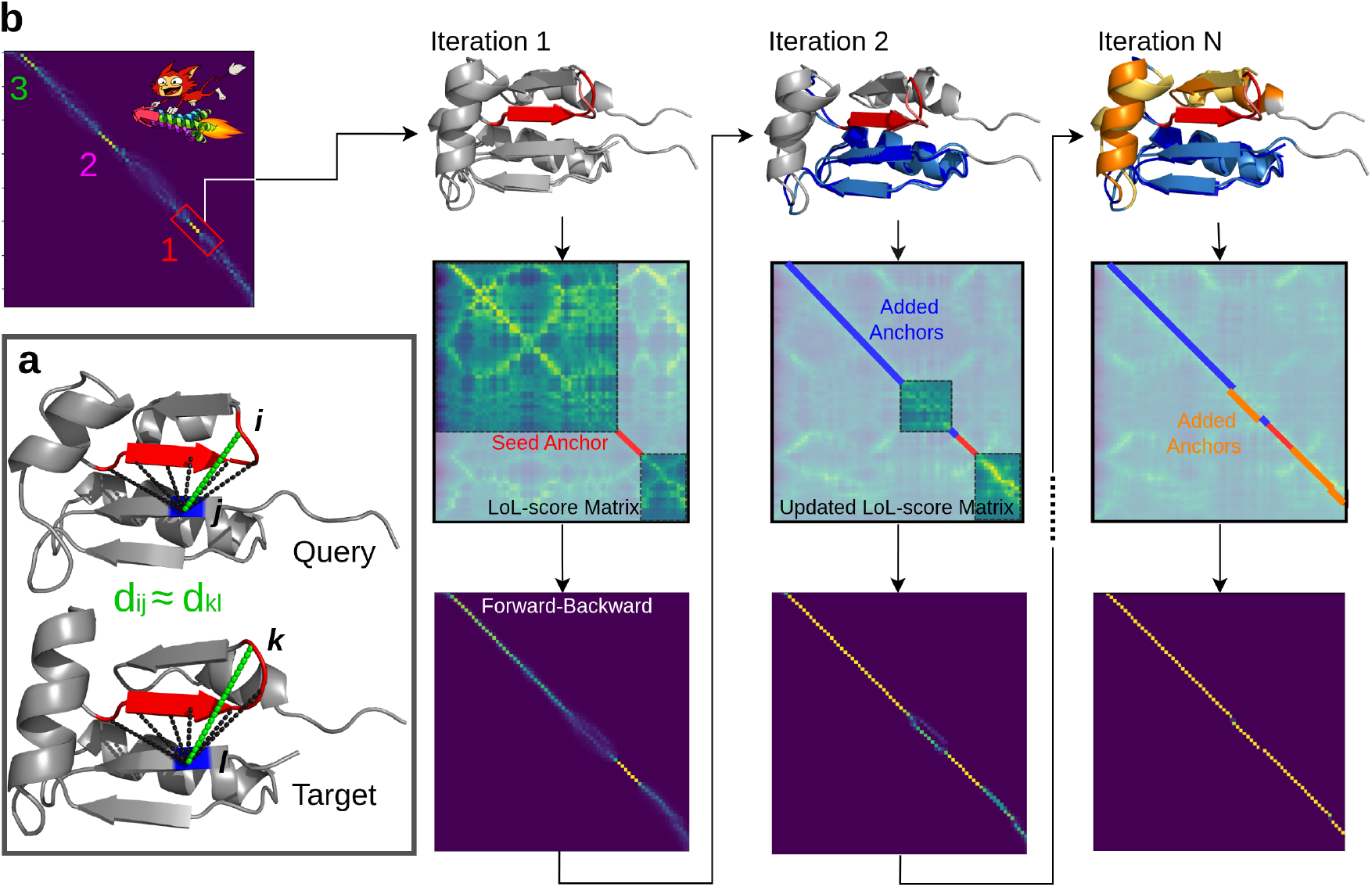
LoL-align algorithm. **a** <u>Lo</u>cal <u>Lo</u>g-odds (LoL) score: A candidate residue pair (*j, l*) (blue) is evaluated against a set *A* of aligned anchor pairs (*i, k*) (red). The LoL-score *S*_*jl*_ quantifies the evidence that *j* and *l* are homologous by comparing intramolecular *C*_*α*_–*C*_*α*_ distances in the query (*d*_*ij*_) and target (*d*_*kl*_) (green): 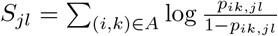, where *p*_*ik,jl*_ is the probability that *i* is homologous to *k* and *j* to *l* given *d*_*ij*_, *d*_*kl*_, and *i*− *j* (see Supplementary Fig. S1). **b.** Algorithm: Top left: Posterior alignment probabilities are computed from 3Di and amino acid substitution scores using our new vectorized Forward–Backward algorithm. Three high-scoring 7-mers are chosen as seed anchors. Top row: Each seed anchor (here 1) is iteratively expanded (colored) until no further residues can be added. Middle row: The LoL-scores are recomputed for all candidate residue pairs with the updated anchor set (colored lines). Bottom row: Posterior probabilities are computed, and pairs whose probability exceeds a threshold are added to the anchor set *A* (top row of next iteration).

In single-domain mode, LoL-align divides the final scores by (*L*_*Q*_ *× L*_*T*_ )^0.25^, while in default multi-domain mode, it uses no normalization. In this mode, if no new anchors are found, the *L*_*Q*_ *× L*_*T*_ 3Di and amino acid score matrices are added to the LoL-score matrix, and the Forward–Backward algorithm is rerun with this combined score. LoL-align can thereby align structures across structural domains even when they fall outside the 20 Å neighborhood of the anchor set. In single-domain mode, this step is omitted to reduce runtime.

We benchmarked the sensitivity of LoL-align and three other popular structural aligners on the SCOPe database [14]. SCOPe classifies single structural domains into a hierarchy of families, superfamilies, and folds. A non-redundant subset, SCOPe40 (11,211 proteins clustered at 40% sequence identity from SCOPe 2.01), was used for evaluation. The query set comprised 3,566 structures, restricted to SCOPe entries with at least one additional family-, superfamily-, or fold-level member. For each query we perform a search against the whole database (11,211) and calculate the sensitivity up to the first false positive, which is the fraction of true positives (TP) found before the first false positive (FP) in the list of ranked matches. TPs are defined to be from the same family, superfamily, or fold, FPs are from different folds.

The cumulative distribution over all queries of sensitivities up to the first FP at the superfamily level in Fig. 2 **a** shows that LoL-align has a 7.3% and 9.9% higher average sensitivity than Dali and TM-align, respectively. In a precision–recall analysis, LoL-align attained an area under the curve 14.5% higher than Dali and 2.3% higher than TM-align (Fig. 2 **b**).

**Figure 2.**
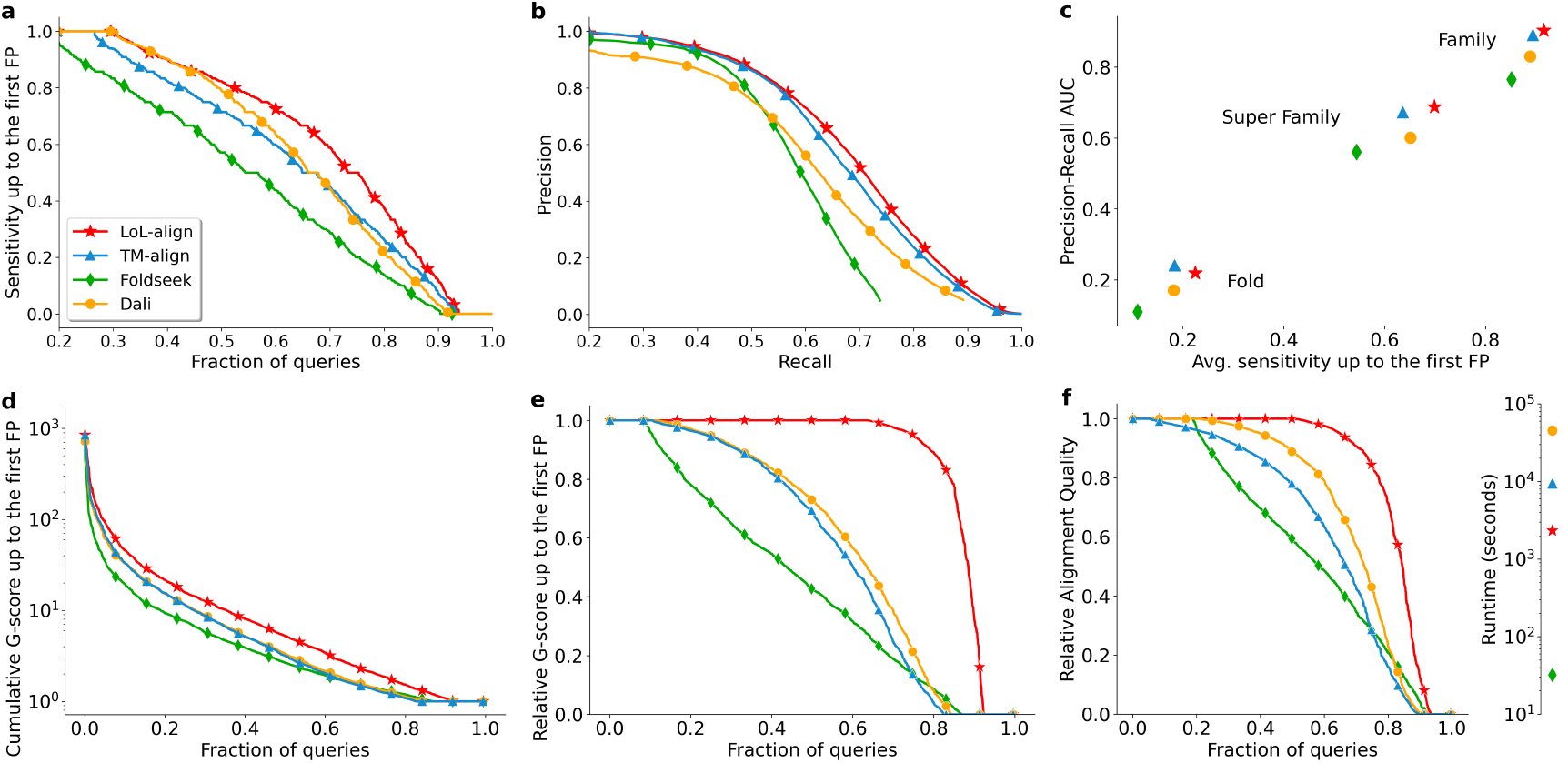
Search sensitivity and alignment quality. **a-c**, homology detection benchmark on the SCOPe40 single-domain structure database. Each structure is searched against all others. True positives (TPs) are matches within the same superfamily, false positives (FPs) are matches between different folds. **a**, Cumulative distributions of the sensitivity for ranking TPs before the first FP. (see Supplementary Fig. S3 for family and fold). **b**, Precision versus recall for superfamily level detection (see Supplementary Fig. S2 for family and fold). **c**, Average sensitivity up to the first FP versus area under the precision-recall curve on family-, superfamily-, and fold-level. **d-e**, Reference-free benchmark on full-length proteins from AFDB. 2000 queries sampled from AFDB were searched against up to 2000 top hits found by Foldseek. **d**, Cumulative rescaled G-scores up to the first FP. The G-score is the geometric mean of residue-wise precision and sensitivity. Alignments with a G-score *<* 0.5 are classified as FPs. G-scores for TPs were rescaled to the interval (0,1). **e**, Relative sensitivity up to the first FP, computed as the cumulative rescaled G-score from panel d normalized by the maximum G-score for the query across all methods. **f**, Cumulative distribution of average alignment qualities, measured as the sum of residue-wise rescaled G-scores for all matches with G-score *>* 0.5 divided by the maximum for the query across all tools. Inset: total runtimes for all 2000 searches.

Searching large databases of full-length protein structures such as the AlphaFold Database [15] (AFDB) is of critical importance nowadays. To assess the alignment quality and search sensitivity on full-length proteins, we selected 2,000 structures from the 2.3 million cluster representatives in the clustered AFDB and sampled 2000 query proteins with at least 100 residues scoring ≥0.7 predicted Local Distance Difference Test (pLDDT) score [1]. Each query was searched against the million structures using Foldseek, retaining up to 2,000 top hits per query as the test set. Each tool was benchmarked by searching each query through these up to 2000 hits.

For each alignment, we calculated its residue-wise precision, defined as LDDT score normalized by the number of aligned residues, and a residue-wise sensitivity, defined as LDDT score normalized by the number of query residues. We calculated the G-score as the geometric mean of precision and sensitivity, and classified alignments as true positives for LDDT G-score ≥0.5 and as true negatives otherwise. (For other cut-offs, see Supplemental Material.) The G-scores were linearly rescaled to the range (0,1) (G-score 0.5 →0, G-score 1.0 →1.0). For each query, we measured search sensitivity as the cumulative rescaled G-score up to the first false positive. LoL-align showed a 40% higher average sensitivity than TM-align and DALI (Fig. 2**d**, note the logarithmic *y* scale). To compare the quality of matches per query on a linear scale, we defined the relative sensitivity for a query as the tool’s search sensitivity divided by the maximum sensitivity observed across all four methods. Again, LoL-align is considerably more sensitive than DALI and TM-align (Fig. 2**e**).

The reference-free nature forced us to assess search sensitivity and alignment quality in a single measure. To assess alignment quality without influence from sensitivity, we reranked all alignments for each query by their G-scores and plotted the cumulative distribution of relative rescaled G-scores (Fig. 2**f** ), demonstrating that LoL-align produces alignments with higher G-scores than the other tools. Finally, on the AFDB benchmark LoL-align was about five times faster than TM-align and twenty times faster than Dali.

In conclusion, LoL-align improves both sensitivity for remote homologs and alignment quality compared with state-of-the-art methods TM-align and Dali, for single domains and full-length proteins. The wide availability of structures for almost any full-length protein sequence calls for tools that combine sensitivity, speed, and alignment accuracy with the ability to detect local similarities and align multi-domain proteins. We demonstrated here that LoL-align addresses these needs by introducing a probabilistic structure similarity score and a new vectorized probabilistic sequence alignment algorithm to efficiently maximize this score. Integration into Foldseek enables systematic functional annotation and evolutionary analyses across hundreds of millions of predicted structures. LoL-align is freely available via the Foldseek web server (search.foldseek.com) or within the open-source Foldseek repo (github.com/steineggerlab/foldseek).

## Online methods

### LoL-align algorithm overview

The LoL-align algorithm is illustrated in Fig. 1. First, the vectorized Forward-Backward algorithm computes for query and target proteins the posterior alignment probabilities for each pair of residues (*i, j*) based on the 3Di sequences. The top ten 7-mer seed anchor candidates with the highest posterior probabilities are re-ranked using gapless alignment centered on the seed using a combination of LoL score, 3Di, and amino acid scores (see Seeds Anchors). The top three candidates are used to initialize full alignments. For each 7-mer seed anchor, the algorithm proceeds iteratively:

1. Compute the matrix of LoL scores using the current anchor residues.
2. Apply the Forward–Backward algorithm to obtain the posterior alignment probability matrix (*P*_*jl*_).
3. Add all residue pairs with *P*_*jl*_ ≥ max_*ik*_(*P*_*ik*_) − 0.1 to the set of anchor residue pairs.

The iteration terminates when no additional anchor pairs are detected. For the multi-domain variant, if no new anchors are found, the 3Di and amino acid score matrices—each of dimension *L*_*Q*_ *× L*_*T*_, where *L*_*Q*_ and *L*_*T*_ denote the lengths of the query and target proteins, respectively—are added to the LoL-score matrix, and the iterative process is resumed from step 2. This allows LoL-align to extend alignments across multiple domains, even when the domains are spatially distant in three-dimensional space.

### The LoL-score

The **Lo**cal **L**og-odds (LoL) score quantifies the evidence that two residues are homologous by evaluating their intramolecular distance relationships relative to other aligned pairs. Let *d*_*ij*_ and *d*_*kl*_ denote the intramolecular *C*_*α*_ distances between residues *i* and *j* in the query and residue *k* and *l* in the target protein. The sequence distance in the query is denoted by *i* − *j*. Let *p*_*ik,jl*_ be the probability that residue *i* is aligned to residue *k* and *j* is aligned to *l*, given the distances *d*_*ij*_, *d*_*kl*_, and *i* − *j*. The residue-wise LoL-score *S*_*jl*_ with respect to a set of anchors *A* is defined as

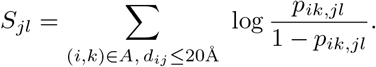

Here, log 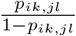 is the log odds that a given residue pair originates from a homologous versus a non-homologous alignment. We approximate this log odds function using a fully connected neural network function *g*(*d*_*ij*_, *d*_*kl*_, *i* − *j*) that takes (*d*_*ij*_, *d*_*kl*_, *i*− *j*) as input, includes a single hidden layer of dimension three with ReLU activation, and outputs a single scalar value. Positive training distance pairs were sampled from TM-align alignments within the same SCOPe superfamily, while negative distance pairs were sampled from alignments between proteins of different folds. The LoL-score for an entire alignment is the sum over the residue-wise LoL-scores for the aligned pairs: LoL-score(*A*) = ∑ _(*j,l*)∈*A*_ *S*_*jl*_.

The LoL-score matrix *S*_*jl*_ does not need to be computed for residue pairs that do not satisfy the strictly monotone alignment constraints (grayed out in Fig. 1). The LoL-scores can be updated incrementally when new anchor pairs *A*_new_ are added, reducing the computational cost of iterative refinement:

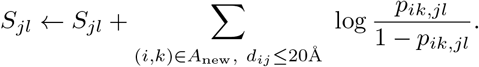

### Selection of new anchor points

Using the LoL scores *S*_*ij*_ derived from the current set of anchor pairs *A*, the Forward–Backward algorithm is applied to compute the posterior alignment probability matrix *P*, where each element *P*_*jl*_ represents the probability that residues *j* and *l* are correctly aligned. New anchor pairs *A*_new_ are then selected as

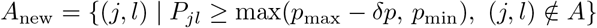

where *p*_max_ = max_*i,k*_(*P*_*ik*_) is the highest posterior probability in *P*, δ*p* is a relative offset from this maximum (default 0.1), and *p*_min_ is the minimum probability threshold for inclusion.

### Vectorized Forward-Backward algorithm

The Forward–Backward algorithm is a widely used dynamic programming method for probabilistic protein sequence alignment. So far, the only implementations we are aware of are sluggish because they are unvectorized and they operate in log space to prevent numerical over- and underflows. The latter requires time-consuming log and exp operations to compute log-sum-of-exp operations. Here, we rewrite the update equations for the Forward-Backward algorithm in a way that allows for vectorization in Python or using the hardware-parallelized SIMD instructions available on modern CPUs. We also demonstrate how to perform the updates in linear space while avoiding under- and overflows.

**The Forward–Backward algorithm** computes from a score matrix *S*_*ij*_ the posterior probabilities *P*_*ij*_ that residue pair (*i, j*) is part of the alignment between a query and a target sequence with lengths *L*_*Q*_ and *L*_*T*_ . Following Mueckstein et al. [13], the probability *P* (*A*) of an alignment *A* is given by *P* (*A*) ∝ *e*^*βS*(*A*)^, with scaling constant *β* and alignment score *S*(*A*) = ∑ _(*ij*)∈*A*_ *S*_*ij*_ − affine gap costs. Normalization to 1 yields the probability distribution

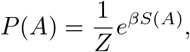

with normalization constant 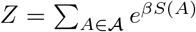 (called partition function *Z*). The sum in *Z* runs over all possible alignments 𝒜.

We define 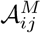as the set of all alignments between residues 1 : *i* from the query and 1 : *j* from the target that end in (*i, j*) ∈ *A* (a “match”). Similarly, we define 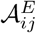 as the set of all alignments between residues 1 : *i* from the query and 1 : *j* from the target that end in a gap in the query protein, and analogously for 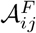 ending with a gap in the target. We then define the partial partition functions 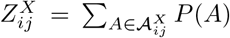 with *X* = *M, E, F* . We denote the gap opening and extension penalties as *g*_o_ and *g*_ext_ (affine gap penalties). It can be shown [13] that these three dynamic programming matrices can be iteratively computed in the forward pass,

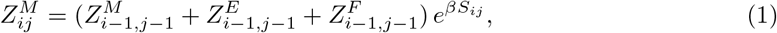

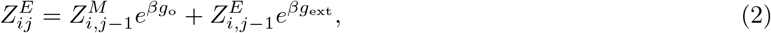

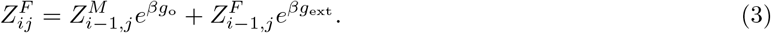

In the backward pass, we compute the matrices 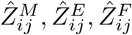 . Here, 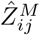 is the sum of *P* (*A*) over all alignments 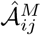 of residues *i* : *L*_*Q*_ in the query and *j* : *L*_*T*_ in the target starting with (*i, j*) ∈ *A*, and analogously for 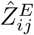 and 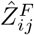 starting with a gap in the query or template, respectively. The update equations for the backward pass are analogous to the ones for the forward pass [13].

Using the matrices from the forward and backward pass, the posterior probability *P*_*ij*_ can be calculated by

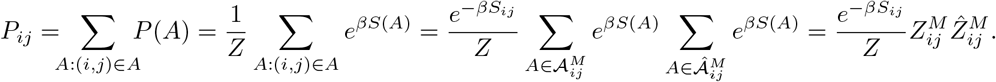

#### Vectorized Forward-Backward

We introduce a new normalization scheme and reformulate key computations to ensure numerical stability in linear space while enabling efficient SIMD vectorization. We divide the target protein into *b* blocks of length *L*. Because the alignment score is additive, the calculation for 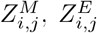, and 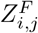 depend only on values from the preceding block (*b* − 1). Each block *b* can therefore be fully processed once the previous block is complete. Within each block, every row *i* is rescaled by max 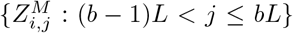 to prevent numerical over- or underflow along the query dimension. These scaling factors are stored in log-space, while all other operations are performed in linear space. To maintain continuity between blocks, we store the final values of 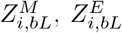, and 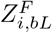 in a compact buffer and use them to initialize the corresponding entries of the next block. These values are updated and then reused as the starting conditions for the next block, ensuring continuity without re-initialization while preventing over- and underflows. The rescaling parameters are incorporated into the final 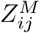 when calculating the posterior probability matrix *P*_*ij*_.

By setting the block length *L* to match the SIMD register width (e.g., *L* = 8 on AVX2 architectures), one operation for a whole line of a block can be processed simultaneously using SIMD vectorization. The recursions for 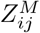 and 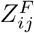 are trivial to vectorize, since they depend only on the preceding row. In contrast, 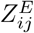 depends on its immediate left neighbor, which prevents direct vectorization. By applying equation (2) iteratively, we obtain

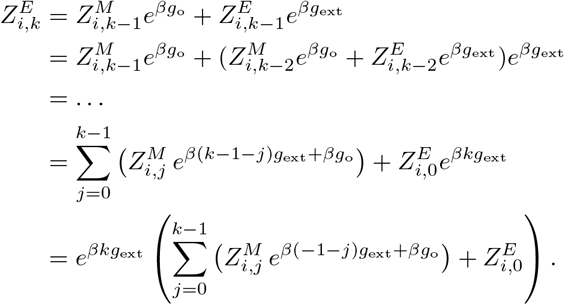

With the precomputed, constant vectors 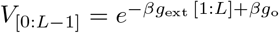 and 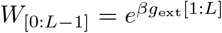, the 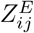 in the first block (*b* = 1) of indices [1:L] can be updated in vectorized fashion,

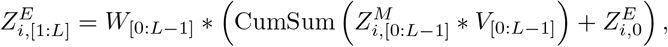

where * denotes element-wise multiplication of vectors. The update equation for later blocks *b* is the same except for an offset (*b* − 1)*L* on all *j* indices.

#### Fast C++ SIMD implementation

To minimize memory usage, a single dynamic programming buffer matrix is reused across all stages of the algorithm. During the forward pass, log-values are calculated from the alignment score matrix and written into the matrix. In the backward pass, these entries are read into SIMD vectors, added with backward contributions in log-space, and written back to the same locations, eliminating the need for a separate backward matrix. Finally, in the posterior calculation step, the matrix is directly aliased as probability matrix *P* and overwritten with posterior probabilities after normalization and lin-space transformation, again in SIMD vectors. All block-level scratch buffers are similarly recycled across passes, so every update happens in-place with vectorized loads and stores, reducing memory traffic and avoiding redundant allocations.

We implemented SIMD-optimized `logf` and `expf` approximations, based on Agner Fog’s Vector Class Library [16], for accelerated evaluation of transcendental functions on 32-bit floats. Both methods apply range reduction followed by minimax polynomial approximation, and compute 4–8 values per instruction using SSE/AVX registers. The `logf` implementation extracts the IEEE-754 mantissa and exponent to evaluate ln(*x*) ≈ *e*· ln 2 + ln(1 + *m*) using an 8th-order polynomial on a normalized mantissa. The `expf` implementation performs argument reduction via log_2_(*e*) and reconstructs the result using a 5th-order polynomial and fast exponent bit synthesis (2^*r*^). The implementations are branch-free aside from masked special-case handling, enabling substantial throughput gains over scalar libm functions while maintaining accuracy suitable for numerically sensitive bioinformatics workloads.

### Seed anchors

The calculation of the LoL-score matrix relies on identifying correctly aligned residue pairs, termed anchors. As seed anchors, we selected high-confidence 7-mers. Anchors were identified using a composite score that combines Foldseek’s 3Di and BLOSUM62 substitution scores, weighted 2.1 and 1.4, respectively, following the Foldseek methodology [10]. Posterior alignment probabilities were computed with the vectorized Forward–Backward algorithm using hyperparameters gap opening = 6, gap extension = 3, and temperature = 2. Anchors were defined as the residue pair with the highest posterior probability together with its three upstream and downstream neighbors. This way, ten non-overlapping 7-mers were chosen as candidate seed anchors and ranked according to the score of the best ungapped alignment passing through their central position. Ungapped alignments were computed using both the composite score (3Di and amino acid) and the LoL-score matrix, with the corresponding 7-mers serving as anchors. The three top-ranked sets were subsequently refined in the iterative alignment process (Fig. 1).

### Score calculation

For ranking alignments, we combine the LoL-score, the 3Di score, and the amino acid (AA) BLO-SUM62 score. To correct for scale differences between residues, we introduce a rescaling factor derived from a self-hit score for each aligned query residue *i*:

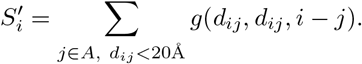

This value represents the maximal LoL-score contribution of residue *i* within an alignment. The rescaling factor is then defined as the mean ratio between observed and optimal contributions:

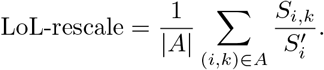

The raw alignment score is multiplied by this factor:

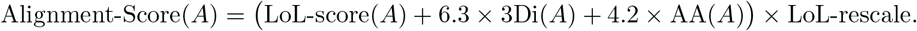

This score is not comparable across queries, as it depends on protein compactness and length. To address this, we normalize by the query–query (QQ) alignment score. In the single-domain version, we additionally normalizes by length by dividing by (*L*_*Q*_ *× L*_*T*_ )^0.25^:

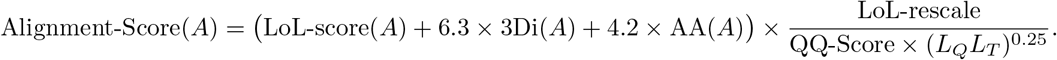

Length normalization improves ranking of single-domain proteins, but can obscure homologous matches in multi-domain proteins. Therefore, in the multi-domain version, the (*L*_*Q*_*L*_*T*_ )^0.25^ factor is omitted.

### Residue-wise sensitivity, precision, and LDDT G-score

The Local Distance Difference Test (LDDT) [17] is a popular measure to assess the quality of protein structure models. It addresses key limitations of older scoring methods such as RMSD, GDT score, and TMscore that relied on global structure superposition, making them not well suited for assessing models of proteins that can undergo conformational changes or of multidomain proteins with poorly defined relative orientations of their domains. LDDT’s recently exploded popularity stems from its use by AlphaFold2 and other deep learning protein structure predictors, which predict together with the protein structure the LDDT score for each residue (predicted LDDT, pLDDT) as a local measure of model accuracy. The LDDT score evaluates each aligned pair of residues (*i, k*) ∈ 𝒜 by a resdiue-wise score between 0 and 1 that describes how well the distances of residues in one structure to its nearest spatial neighbor residue *j* agree with the distances bewteen the aligned residues in the aligned (model) structure. Each pair of homologous distances (*d*_*ij*_, *d*_*kl*_) obtains a residue-wise score between 0 and 1, 1 for very good agreement (*d* = *d*_*ij*_ − *d*_*kl*_ *<* 0.5Å), 0.75, 0.5, and 0.25 for partial agreement, and 0 for a deviation of more than 4Å. The residue-wise score for residue pair (*i, k*) is the mean of these *N*_*i*_ distance scores. The LDDT score is the sum of residue-wise scores for all aligned pairs divided by the number of query residues *L*_*q*_:

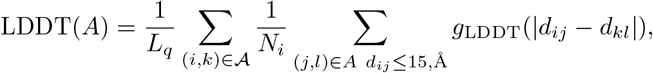

where 𝒜 is the set of aligned residues, *N*_*i*_ is the number of aligned neighbors within 15 Å, and, *g*_LDDT_ is a step function that scores the difference *d* between two intramolecular distances,

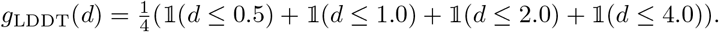

The LDDT score thus answers the question what fraction of query residues was aligned correctly to the target structure, while considering a continuous measure for correctness in the form of the residue-wise LDDT score. When using it as a score for the quality of a structural alignment, it therefore measures residue-wise sensitivity (a.k.a. recall): What fraction of aligned query residues was actually correctly aligned. Taken alone, it favors overprediction of aligned residue pairs because it does not penalize overpredictions.

To address this, we could normalize the LDDT score by the number of aligned residues |𝒜| instead of the number of query residues. This score measures the fraction of aligned query residues that was actually correctly aligned, which is the residue-wise precision. Taken alone, it favors underprediction, omiting all but the most reliable parts of the alignmemt.

Neither sensitivity nor precision alone are a good measure of performance since we need both to be high for an alignment to be useful. A common balanced measure is the F1 score, equal to the harmonic mean of sensitivity and precision. However, since our precision is always greater than or equal to sensitivity, the F1-score can disproportionately emphasize sensitivity. We therefore chose the G-score, the geometric mean of precision and sensitivity.

When working with protein structures predicted by AlphaFold, mispredicted regions must be handled carefully. Residues in such regions typically have low pLDDT scores. Therefore, we used only residues with pLDDT ≥0.5 to calculate the LDDT score and adjusted both the query and alignment lengths accordingly when normalizing the score.

### Full length Benchmark

To assess performance on full-length proteins, we sampled 2,000 cluster representatives from the clustered AlphaFold database [15], selecting only proteins with at least 100 residues scoring ≥0.7 pLDDT. Each query was searched against all representatives using Foldseek (parameters: `-s 9.5 -e 10 --max-seqs 2000`). The resulting hits were aligned with Dali, TM-align, LoL-align, and Foldseek, and ranked by their respective scores. For TM-align, we used the TM-score normalized by query length, which performed best on this benchmark. LoL-align was used in the multi-domain version (`--lolalign-multidomain 1`).

For each alignment, we calculated the residue-wise precision and sensitivity. Alignments with an LDDT G-score ≥0.5 were classified as true positives (see Supplementary Fig. S4 for alternative thresholds). To give very good alignments more weight than marginally correct ones, G-scores were linearly rescaled from (0.5,1) to the range (0,1) via Score = 2 (G-score − 0.5). We measured search sensitivity as the cumulative rescaled G-score up to the first false positive, including self-hits to avoid zero scores in log space (Fig. 2**d**). We then defined relative sensitivity as this search sensitivity divided by the maximum sensitivity across all methods for that query, here we exclude self-hits (Fig. 2**e**). Finally, to assess alignment quality independently of method-specific ranking, we ranked all alignments by their G-scores (excluding self-hits) and plotted the cumulative distribution of relative rescaled G-scores (Fig. 2**f** ).

We note that LoL-align and Dali optimize scores more closely related to the LDDT metric, as all are based on comparing intra-molecular distances, in contrast to TM-align. We selected the LDDT-based G-score because metrics requiring structural superposition, such as TM-score or RMSD, do not adequately capture alignment quality for full-length, multi-domain proteins and larger proteins with conformational flexibility, which were the focus of this reference-free benchmark.

All alignment tools were executed on a dual-socket AMD EPYC 7742 system with 128 physical cores. Tools with native multithreading support (LoL-align and Foldseek) were run with 128 threads. Tools without multithreading support were parallelized by splitting the query set into 128 equally sized batches and processing them in parallel.

### SCOPe Benchmark

For the SCOPe benchmark, alignment results for Dali, TM-align and Foldseek were reused from the Foldseek study [10], where the details of the alignment procedures are described. LoL-align was applied in the single-domain setting (`--lolalign-multidomain 0`).

### Evaluation SCOPe benchmark

After sorting the alignment results for each query, sensitivity was calculated as the fraction of true positives (TPs) in the ranked list up to the first false positive (FP), excluding self-hits. Mean sensitivity was compared across all queries at the family, superfamily, and fold levels. Only SCOPe entries with at least one additional family-, superfamily-, or fold-level member were included. Sensitivity was also measured up to the first FP (ROC1), rather than to the *k*-th FP (e.g., the 5th FP), as ROC1 better reflects the requirements of low false discovery rates in automatic searches.

Precision–recall curves were generated for each tool. Alignment results were sorted by structural similarity scores, and self-matches were excluded. Curves were computed separately at the family, superfamily, and fold levels, with precision defined as TP*/*(TP +FP) and recall as TP*/*(TP +FN). All counts (TP, FP, FN) were weighted by the reciprocal of their respective family, superfamily, or fold size, ensuring that families, superfamilies, and folds contributed linearly with their size rather than quadratically.

### LoL-align usage

LoL-align can be used within Foldseek’s easy-search by setting `--alignment-type 3`, which runs Foldseek’s search (3Di prefilter and 3Di+AA alignment) and realigns the top hits using LoL-align. To use LoL-align without Foldseek’s prefilter, a fake prefilter can be created as described in the [MMseqs2 user guide](https://mmseqs.com/latest/userguide.pdf), and the alignment can then be executed with `foldseek lolalign queryDB targetDB prefilter results`.

On the Foldseek server, LoL-align can be used to realign search hits by selecting the LoL-align option under Databases & search settings.

## Acknowledgments

The following support is gratefully acknowledged: J.S. by the German Research foundation (DFG) (RTG2984 Evorest), M.S. by the National Research Foundation of Korea (RS-2020-NR049543, RS-2021-NR061659 and RS-2021-NR056571, RS-2024-00396026), Creative-Pioneering Researchers Program and Novo Nordisk Foundation (NNF24SA0092560), and M.M. by the National Research Foundation of Korea (grant RS-2023-00250470).

## Author contributions

**L.R**. Designed the research; Developed the code; Performed analyses; Writing - Original Draft; Writing - Review & Editing. **M.v.K**. Designed the research; Developed the LoL-Score; Writing - Review & Editing. **G.K**. Developed the code; Writing - Review & Editing. **S.H.K**. Developed the code; Writing - Review & Editing. **M.M**. Developed the code; Integrated LoL-align into the Foldseek server; Writing - Review & Editing. **M.R**. Performed analyses. **M.S**. Developed the code; Writing - Review & Editing. **J.S**. Designed the research; Developed the LoL-Score; performed analyses; Writing - Original Draft; Writing - Review & Editing.

## Data availability

Data utilized to perform benchmarks for this study are freely available at https://wwwuser.gwdguser.de/compbiol/lolalign/

## Code availability

All code developed in this study is available under GPLv3 license and documented at foldseek.com. The SIMD Forward-Backward algorithm is implemented in MMseqs2 under MIT license at mm-seqs.com. Analysis scripts are available at github.com/soedinglab/LoLalign-analysis.

## Supplementary information

**Figure S1:**
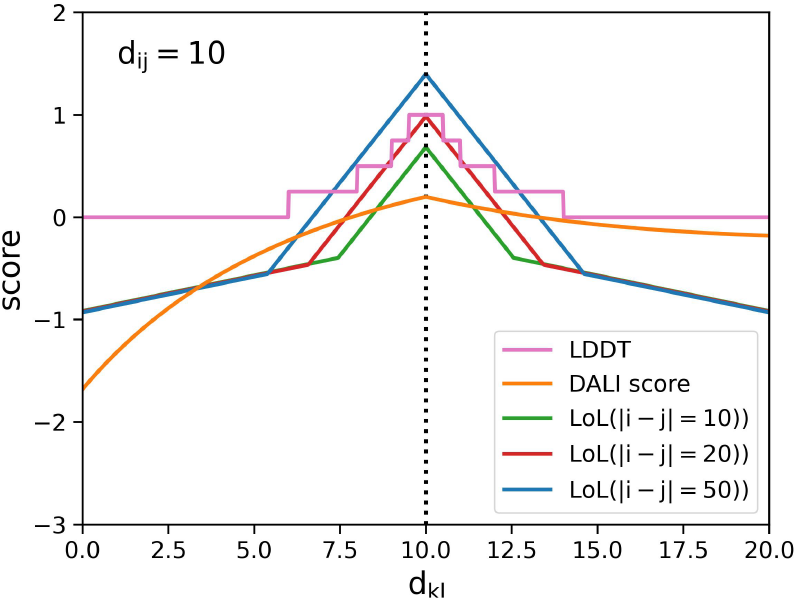
Visualization of the LoL-Score. Visualization of the LoL-score with different sequence distances as well as the Dali- and LDDT-score.

**Figure S2:**
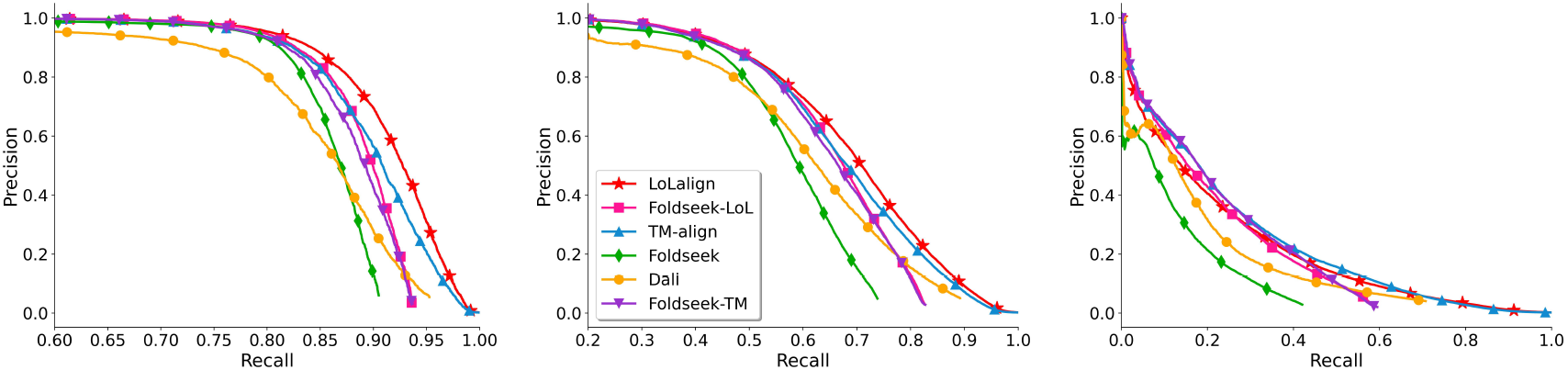
Precision-Recall curves on SCOPe benchmark. We measured the sensitivity on the SCOPe40 dataset by plotting precision-recall curves on family, superfamily, and fold level. Where Foldseek-TM and Foldseek-LoL uses the Foldseek search with `-s 9.5 -e 10 --max-seqs` 4000 as a prefilter.

**Figure S3:**
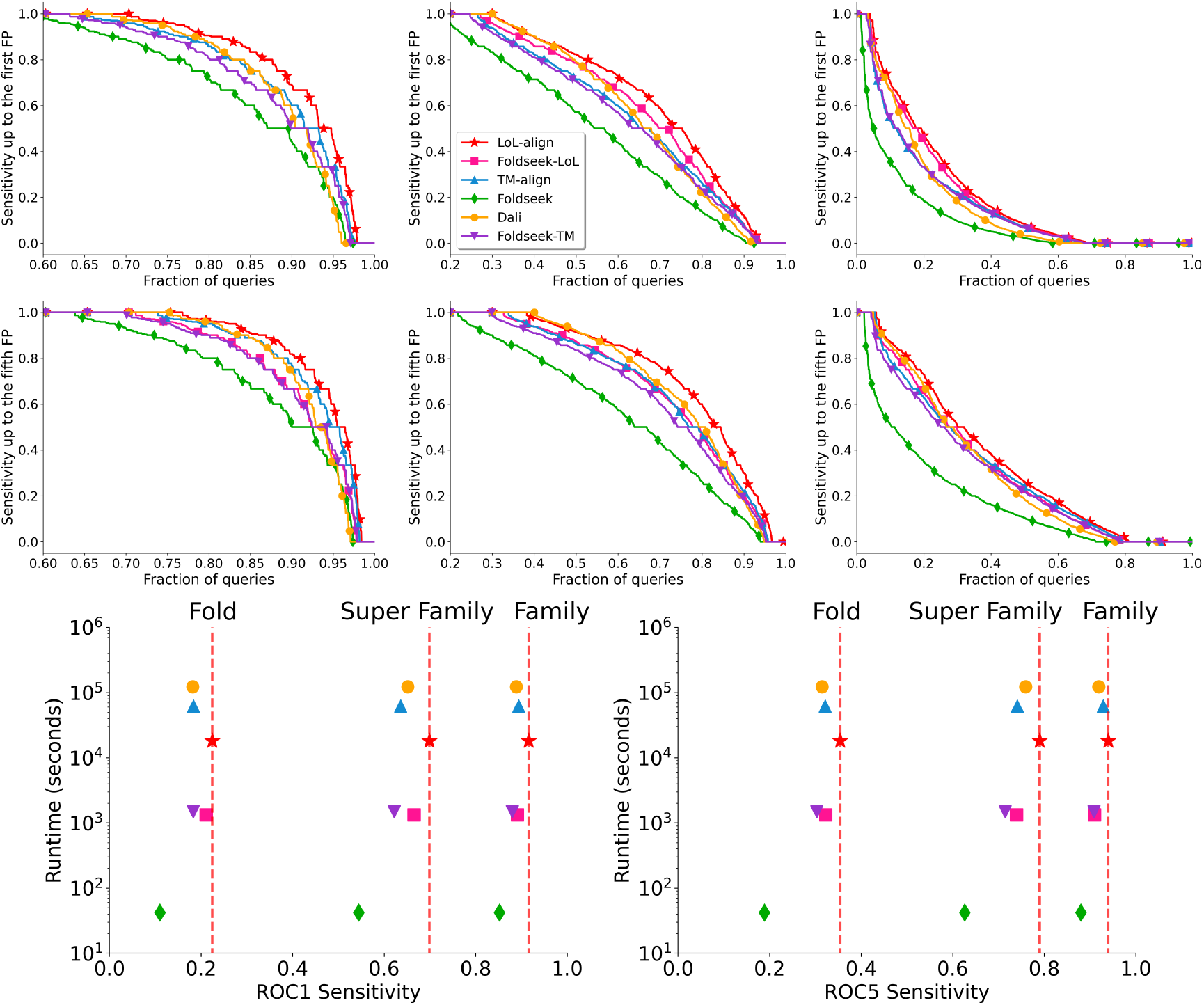
cumulative distributions of sensitivity for the SCOPe benchmark. We measured the sensitivity on the SCOPe40 dataset by plotting the cumulative distributions of sensitivity up to the first and fifth FP on family, superfamily, and fold level. Where Foldseek-TM and Foldseek-LoL use the Foldseek search with `-s 9.5 -e 10 --max-seqs` 4000 as a prefilter. We also show the average ROC1 and ROC5 sensitivity plotted against the runtime. For TM-align we used the -fast parameter.

**Figure S4:**
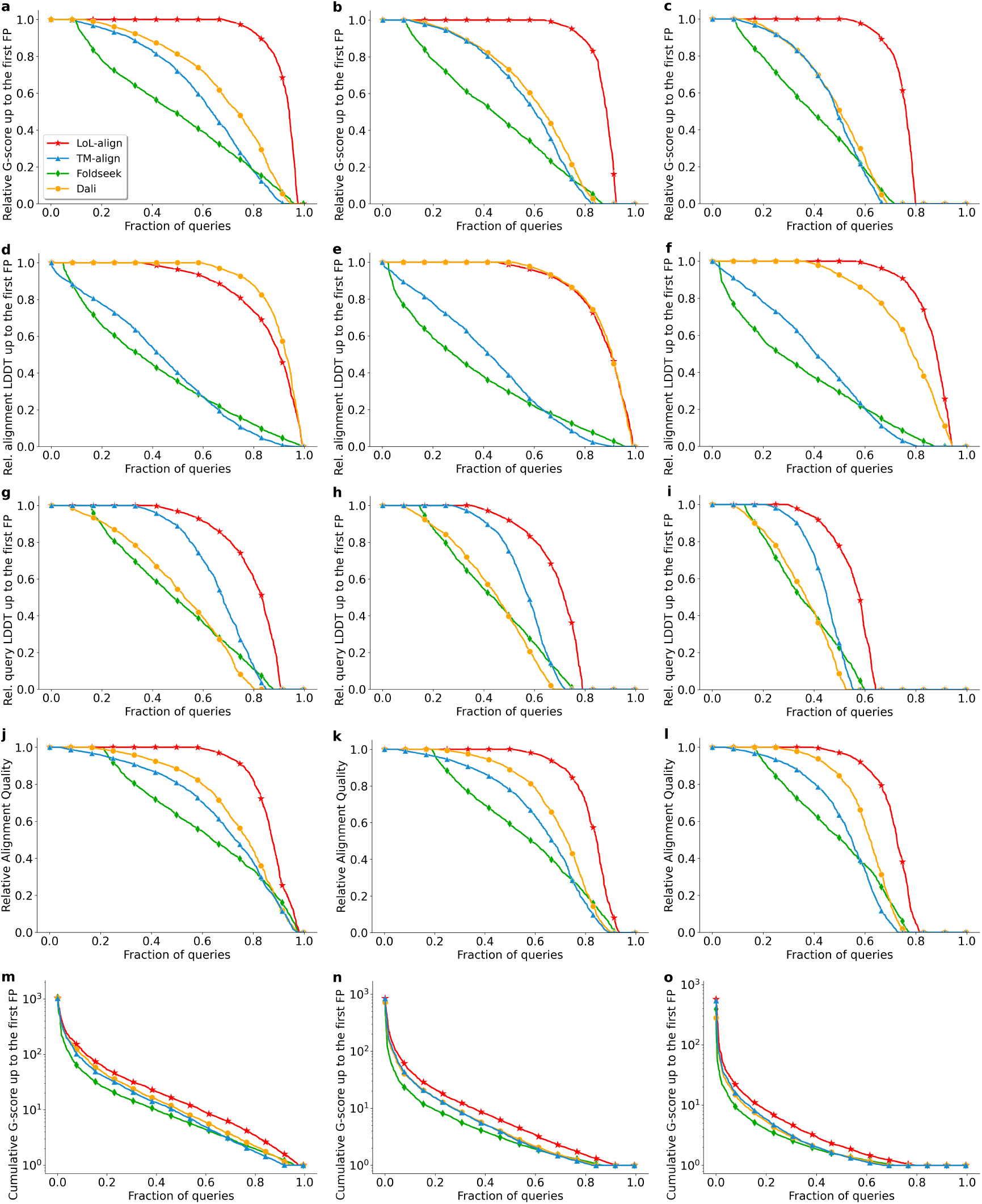
Detailed analysis of the full-length structure alignment benchmark. Each column corresponds to a different cutoff used to define true positives (0.4, 0.5, and 0.6 from left to right). The first row (**a–c**) shows the same benchmark as in Fig. 2e. The second row (**d–f** ) displays results for the LDDT score normalized by alignment length (precision score). The third row (**g–i**) shows results for the LDDT score normalized by query length (sensitivity score). The fourth row (**j–l**) reproduces the analysis from Fig. **2f**, and the fifth row (**m–o**) corresponds to the analysis shown in Fig. **2d**.

